# Quantification of autism recurrence risk by direct assessment of paternal sperm mosaicism

**DOI:** 10.1101/208165

**Authors:** Martin W. Breuss, Morgan Kleiber, Renee D. George, Danny Antaki, Kiely N. James, Laurel L. Ball, Oanh Hong, Camila A. B. Garcia, Damir Musaev, An Nguyen, Jennifer McEvoy-Venneri, Renatta Knox, Evan Sticca, Orrin Devinsky, Melissa Gymrek, Jonathan Sebat, Joseph G. Gleeson

## Abstract

*De novo* genetic mutations represent a major contributor to pediatric disease, including autism spectrum disorders (ASD), congenital heart disease, and muscular dystrophies^1,2^, but there are currently no methods to prevent or predict them. These mutations are classically thought to occur either at low levels in progenitor cells or at the time of fertilization^1,3^ and are often assigned a low risk of recurrence in siblings^4,5^. Here, we directly assess the presence of *de novo* mutations in paternal sperm and discover abundant, germline-restricted mosaicism. From a cohort of ASD cases, employing single molecule genotyping, we found that four out of 14 fathers were germline mosaic for a putatively causative mutation transmitted to the affected child. Three of these were enriched or exclusively present in sperm at high allelic fractions (AF; 7-15%); and one was recurrently transmitted to two additional affected children, representing clinically actionable information. Germline mosaicism was further assessed by deep (>90x) whole genome sequencing of four paternal sperm samples, which detected 12/355 transmitted *de novo* single nucleotide variants that were mosaic above 2% AF, and more than two dozen additional, non-transmitted mosaic variants in paternal sperm. Our results demonstrate that germline mosaicism is an underestimated phenomenon, which has important implications for clinical practice and in understanding the basis of human disease. Genetic analysis of sperm can assess individualized recurrence risk following the birth of a child with a *de novo* disease, as well as the risk in any male planning to have children.

## Main Text

A newborn child harbors, on average, 40-80 *de novo* single nucleotide variants (dSNVs) across the genome^1,3^, which have the potential to influence health of the child and impact biological fitness^6^. Consequently, severe early-onset conditions, such as congenital heart disease, epilepsy, or intellectual disabilities are enriched for *de novo* mutations that activate or inactivate critical genes^7^. Although this applies broadly to autism spectrum disorders (ASD), high impact dSNVs are concentrated among a subset of cases with low IQ^8^.

The majority of *de novo* mutations originate in the parental germline^9–11^. Depending on the timing and location of the mutation, these may be mosaic throughout the body or only in the parental germline, and they could transmit to multiple offspring. Indeed, this has been documented in families with recurrent *de novo* mutations as well as in recent studies that assessed parental mosaicism^1,9,10^. Two crucial questions, however, remain unaddressed: What is the extent of mosaicism in the parental germline? Can quantification of mosaicism help to estimate risk of recurrence in the parents?

We propose three main hypotheses: 1) a subset of dSNVs originate from early mutational events and can be detected as mosaic in the germ cell population; 2) due to the early separation of the germline during embryogenesis, these dSNVs are either absent or underrepresented in peripheral tissues; and 3) by assessing germ cells directly, prediction of the recurrence risk of a given mutation may be accurately assessed.

Because a majority of dSNVs are paternal in origin^9,11^, and because germline allele frequencies can be quantified directly from sperm-derived DNA, we focused on the profiling of germline mosaicism in fathers of children affected with ASD. We accessed genetic data from two independent ASD cohorts: one focusing on 98 probands with ASD and an additional diagnosis of epilepsy (O.D. and J.G.G; unpublished), and one consisting of 71 probands with general features of ASD (J.S.)^12^. For both, candidate dSNVs were identified through unbiased trio *de novo* sequencing^8,13^. Twelve families met criteria, where a likely pathogenic mutation was identified, and where participants agreed to provide semen samples (Fig. 1a and Supplementary Table 1). We used single molecule droplet digital PCR (ddPCR) genotyping for quantitative analysis of mosaicism, with a detection limit of 0.1% allelic fraction (AF) (Fig. 1a).

**Figure 1.**
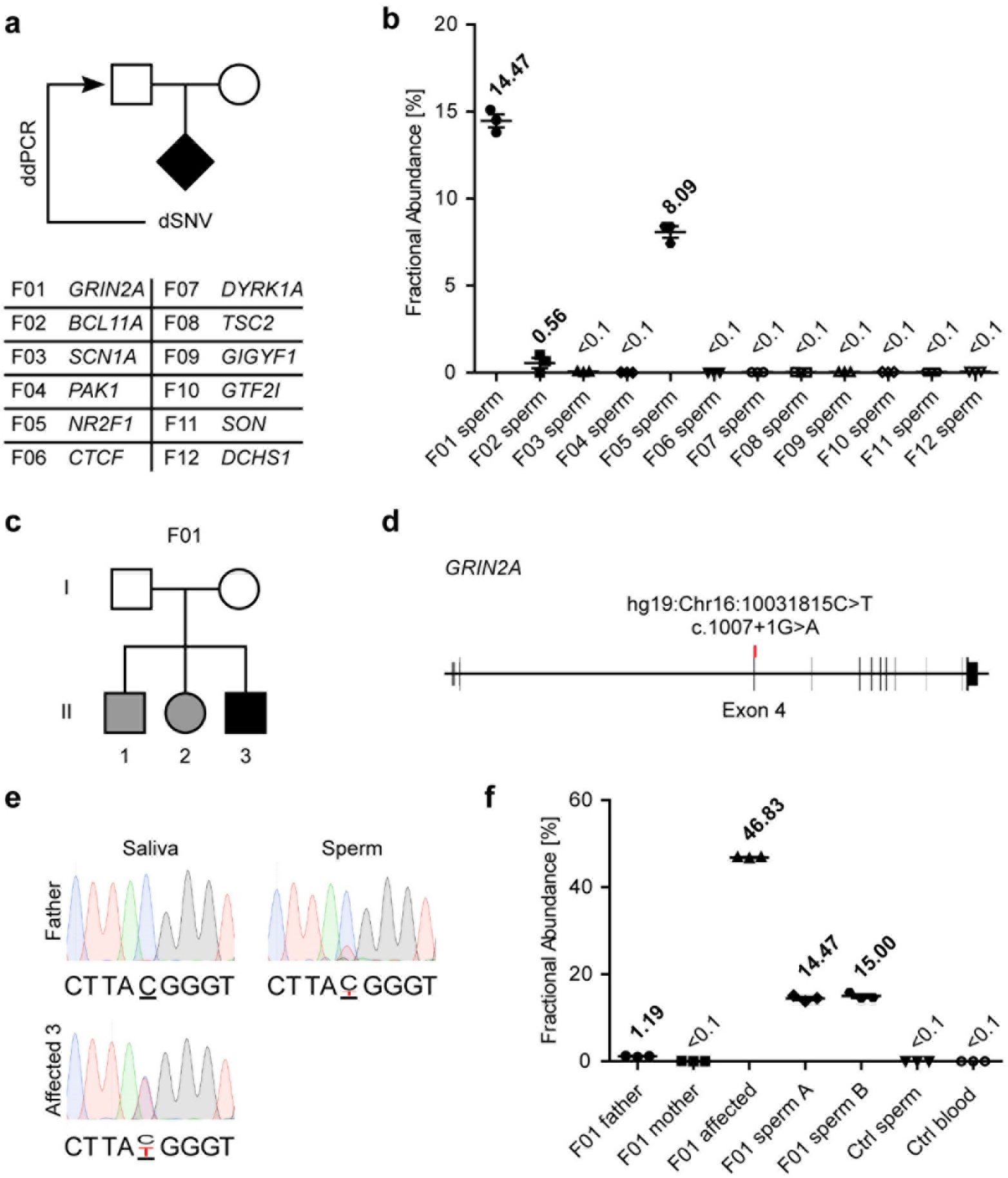
Inherited, pathogenic *de novo* SNVs are detected in paternal sperm in three out of ASD cases. **a**, Schematic of the ASD trios that harbor *de novo* single nucleotide variants (dSNVs) and a list of the interrogated genes in the 12 families. The likely pathogenic dSNVs were quantified in the paternal sperm by ddPCR. **b**, Fractional abundance (determined by ddPCR) of the mutant allele in paternal sperm for the relevant dSNV for the 12 families. Thisdata along with control reactions on unrelated blood and sperm is also shown in Extended Data Fig. 1 a,b. **c**, Pedigree of F01 showing the parents in generation I and three affected children in generation II. Black indicates ASD, grey, epilepsy with ADHD symptoms (see Supplementary Table 2). Note that only II-3 met the criteria for *bona fide* ASD and was the only child included in the original trio-based study. **d**, Schematic of *GRIN2A* and the mutation that was found in all three children and affected a splice site. **e**, Sanger sequencing results showing the C>T conversion described in **d**. The affected child was heterozygous for the mutation and paternal sperm showed a minor peak of the mutant allele consistent with the ddPCR results. **f**, Fractional abundance plot as in **b** for the *GRIN2A* mutation in the F01 family. Father, mother, and affected indicate saliva samples of the respective individual, whereas sperm A and B indicate biological replicates of the paternal sperm. Ctrl-an unrelated sperm or blood sample, as indicated, used as control. Plots in **b** and **f** show the individual data points (technical triplicates), as well as the mean ± SEM.

Of the 12 samples, three harbored mosaic variants in father’s sperm (Fig. 1b, Extended Data Fig. 1a-c and 2a-c), two of which showed clinically significant AF of 14.47% (F01) and 8.09% (F05). This contrasts with the *a priori* risk in the general population of 1:68 (i.e. ~1.5%) for ASD, suggesting a 5-10 fold elevated risk for these males of fathering a subsequent child with the same mutation. Strikingly, the matched blood or saliva exhibited either a drastically lower fractional abundance (1.19%, F01) or were below the detection limit of our assay (<0.1%, F05) (Fig. 1f and Extended Data Fig. 2b). The variant found in F02 showed similar mosaicism in sperm (0.56%) and saliva (1.16%) (Extended Data Fig. 2a). However, low sperm counts and amount of recovered DNA may render this sample sensitive to artifacts (e.g. somatic tissue contamination or biased sampling of a small subpopulation of cells).

Our original trio-analysis of F01 only included one affected child (II-3), harboring a *de novo* mutation in *GRIN2A*, a cause of diverse neurodevelopmental conditions (Fig. 1c,d)^14,15^. Following disclosure of the mosaic germline variant, the parents reported that both older children displayed neurological phenotypes consistent with this mutation (Fig. 1c and Supplementary Table 2)14,15. Indeed, we found that both carried the same heterozygous, pathogenic variant that was detected in the proband and mosaic in the father (Fig. 1e,f and Extended Data Fig. 1d), although the expressivity in this family argued against a single genetic cause (Supplementary Table 2). (Only the epilepsy, but not ASD was expressed in all children. The youngest sibling met criteria for ASD, whereas the middle child showed features of ADHD and speech impairment; the oldest sibling had ADHD, which resolved in adolescence.) This initial analysis of a limited ASD cohort confirmed our three hypotheses and argued that paternal sperm can provide useful material for genetic assessment, which can impact diagnosis and discussions of recurrence risks.

Having assessed only a single variant in each family, we were not able to draw conclusions regarding the full extent of germline mosaicism. Therefore, we additionally performed whole genome sequencing (WGS) of matched sperm and blood samples at >90x read depth from four fathers (F08, F09, F21, and F22) (Extended Data Fig. 3a,b). We first assessed paternal sperm mosaicism for 355 high-confidence dSNVs detected in their six children (denoted ‘all’, Fig. 2a). We confirmed that the number per offspring increased as a function of the paternal age at conception (R^2^=0.804, P=0.015) (Fig. 2b). This was consistent with previous reports and provided evidence that these dSNVs represented biologically relevant mutational events^9,11^. From these, 152 variants were phased to the paternal haplotype (denoted ‘pat’, Fig. 2a) and also showed the expected positive correlation with paternal age (R^2^=0.939, P=0.031) (Fig. 2b). In the paternal WGS data, 16.62% (all) and 19.08% (pat) of these dSNVs were detected in the father, the majority being exclusively present in sperm, or enriched in sperm relative to blood (Fig. 2c and Supplementary Table 3). Most variants were present at AF below 2% and were uniformly distributed across chromosomes (Fig. 2d,e and Extended Data Fig. 4a-d,g). Neither the number of mosaic dSNVs, nor the average AF were significantly correlated with the paternal age at conception (Extended Data Fig. 4e,f).

**Figure 2.**
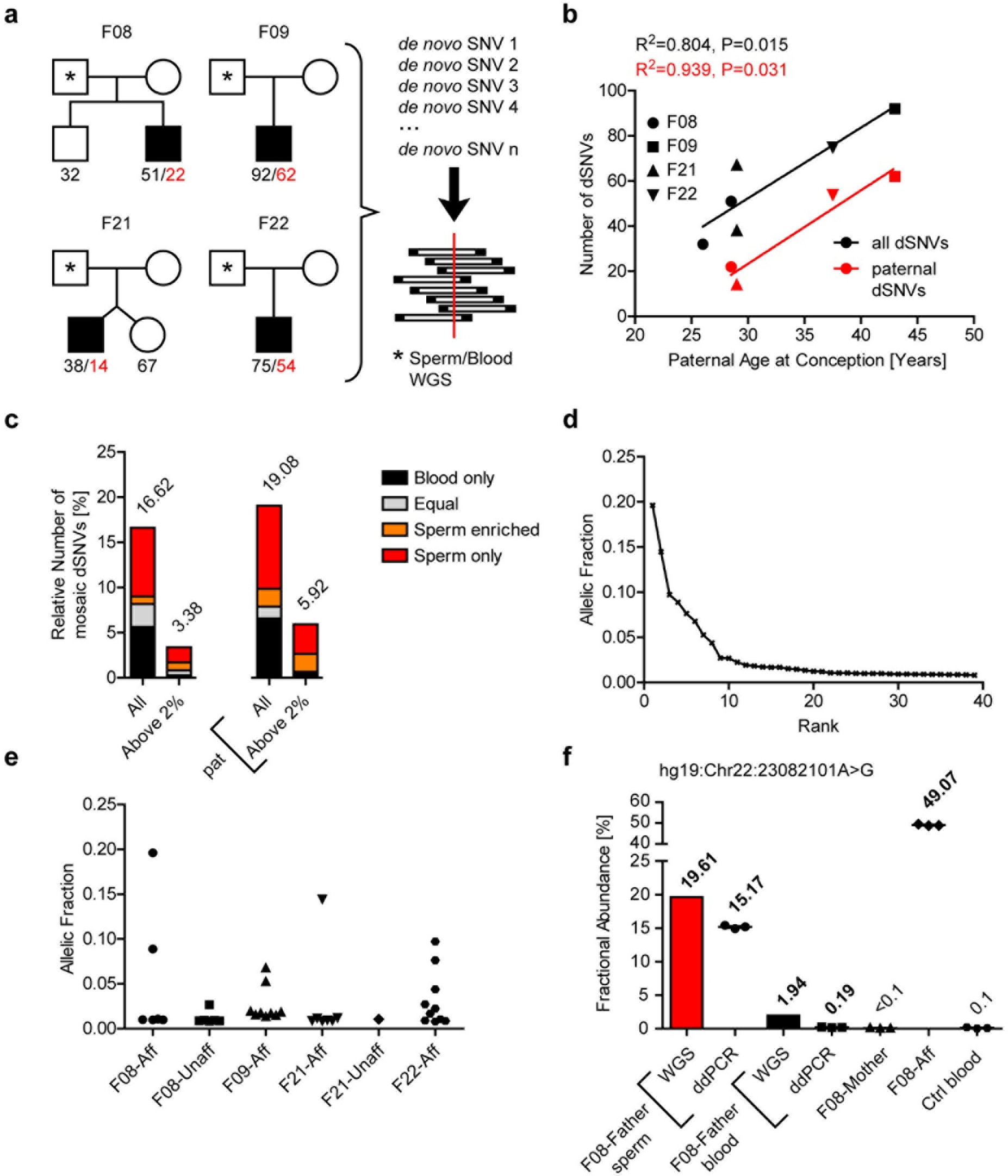
Frequencies and allelic fractions of dSNVs in paternal sperm. **a**, Schematic showing all four pedigrees for F08, F09, F21, and F22 and the number of detected dSNVs for each child. Black numbers indicate all dSNVs that were detected in a given individual, whereas red numbers indicate the subset of variants from the affected that were phased to the paternal haplotype using Oxford Nanopore long-read (average read length: 6,772 bp) technology. The right side depicts the basic strategy: detected dSNVs were assessed in the respective father’s sperm and blood using WGS data. **b**, Plot showing the increase in dSNV number with paternal age at conception, as expected^9,11^. Line shows a regression curve demonstrating this dependence (all dSNVs: R^2^=0.804, P=0.015; paternal dSNVs: R^2^=0.939, P=0.031). Red font indicates data for those dSNVs that were phased to the paternal haplotype. **c**, Quantification of the relative number of dSNVs that showed evidence of mosaicism in blood, sperm, or both. Equal denotes variants that were detected at roughly equal ratios in both data sets (α<3). All: relative numbers of dSNVs at all AF; Above 2%: relative numbers of dSNVs at AF>2%; pat: data from paternally phased dSNVs only. The results show that most variants above 2% were either found only in sperm or were enriched in sperm **d**, Plot of all mosaic variants detected in sperm versus respective allelic fraction (AF). **e**, Same data set as in **d**, but separated by which child harbored the dSNV. **f**, Fractional abundance (determined by ddPCR or WGS read counts) of the mutant allele Chr22:23082101A>G in family F08. Mother and Aff depict blood samples from the mother and the affected child that harbored this mutation. Graph shows individual data points (technical triplicates) and mean ± SEM for the ddPCR data.

Orthogonal validation by Sanger sequencing and ddPCR confirmed germline presence of most variants with AF above 2% (5/6 by Sanger sequencing), but not those below this threshold (0/6 by ddPCR) (Fig. 2f and Extended Data Fig. 5). Based on this and the baseline risk of ASD in the general population of ~1.5%, we used 2% as the threshold for clinically relevant mosaicism. Of all dSNVs, 3.38% (all) and 5.92% (pat) exceeded this level (Fig. 2c). These data suggest that, in the absence of selection acting on the pathogenic mutation, more than 3% of diseases caused by dSNVs have a risk of recurrence that is substantially elevated when compared to the basal population-wide risk^16^. Moreover, for the majority of cases this risk cannot be assessed accurately in somatic tissues, but can be assessed in paternal sperm.

We also studied the genome-wide extent of mosaic variation in sperm using the WGS data (Fig. 3a). We identified high-quality sperm-specific mosaic variants using a stringent pipeline that utilized two mosaic detection algorithms: MuTect and Strelka. We further excluded likely false calls in repetitive regions or within ±5 base pairs of a germline insertion or deletion variant. In total, we detected 30 mosaic SNVs with AF between 29.8% and 3.7% in the sequenced sperm from the four fathers (Fig. 3b-e, Extended Data Fig. 6a-d, and Supplementary Table 4). As with mosaic dSNVs, neither the number of mosaic SNVs, nor their average AF showed correlation with age at sampling (Extended Data Fig. 6e-f).

**Figure 3.**
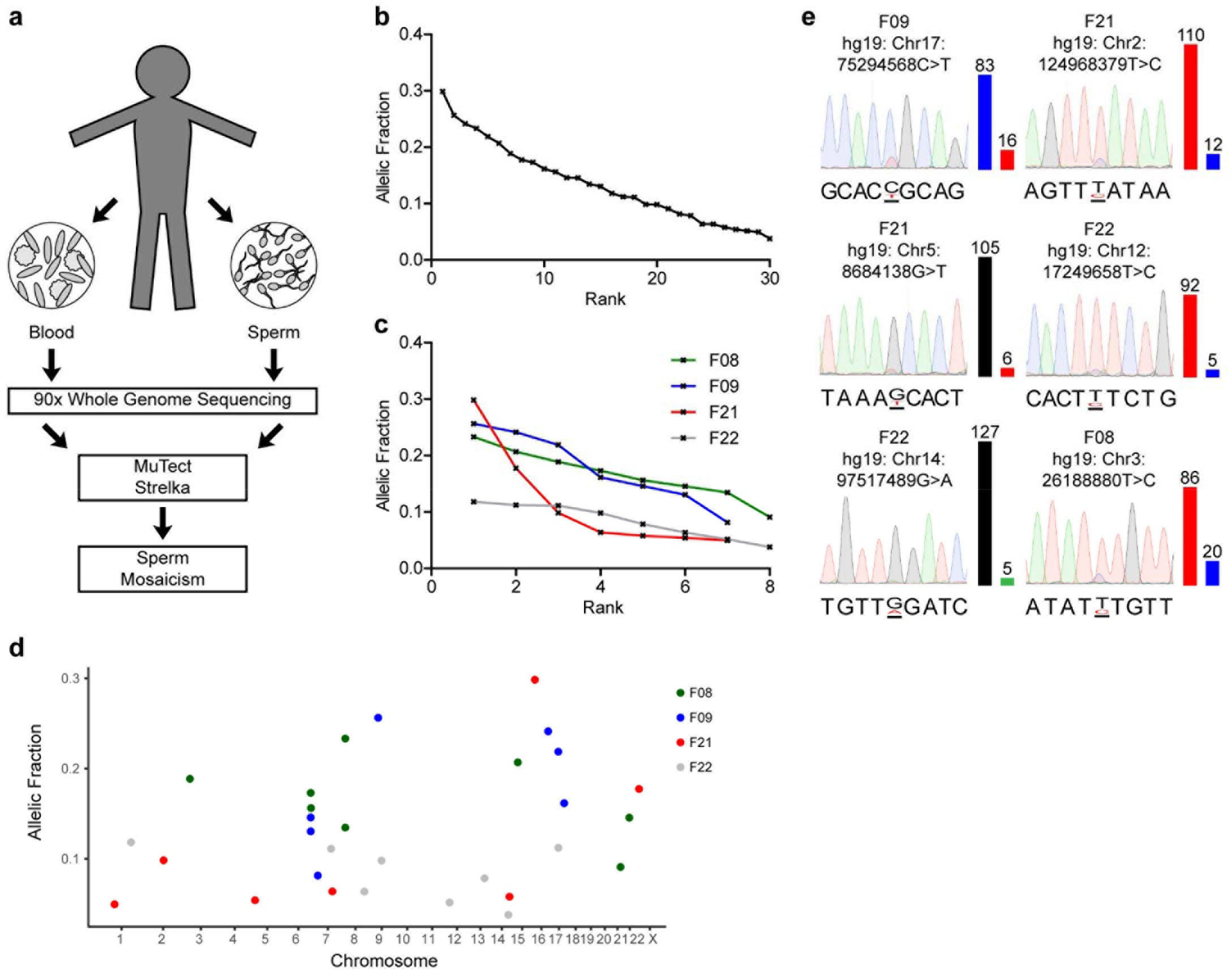
Unbiased detection of mosaic variants in sperm. **a**, Schematic illustrating the workflow for the analysis of mosaicism in sperm. **b**, Plot of all mosaic variants detected in sperm and respective AF. **c**, Same data set as in **b**, but separated by family. **d**, Plot of all mosaic variants showing roughly equal distribution across the genome. **e**, Sanger sequencing results for six of the detected mosaic variants, with relative peak height representing degree of mosaicism. Beside the chromatograms, bars depict read counts in the WGS data for the reference allele onthe left and the mutant allele on the right and are colored according to the base they represent (A: green, T: red, G: black, C: blue).

We likely underestimated the true number of mosaic SNVs, as we could only detect mosaic variants above 4% AF at current sequencing depth of >90x. Moreover, we only detected the top-ranked variant (Chr22:23082101A>G in F08) from the pool of 39 mosaic dSNVs (Fig. 2f and Supplementary Table 4). Future efforts to detect lower frequency, pathogenic variants in sperm will require improved algorithms for detection, higher sequencing depths, or both. A cost-effective way to achieve this would be to sequence targeted regions of interest (e.g. the exome, haploinsufficient genes, or candidate genes for sporadic diseases). Nevertheless, the relatively high frequency of germline mosaicism with variants present at high AF argues for the clinical utility of screening sperm to identify carriers prior to conception. Furthermore, a complementary analysis of blood-specific mosaicism suggested a largely distinct set of variants from those evident in the germ cells (Extended Data Fig. 7). This should be explored further by increasing the sensitivity of mosaic detection, which would allow for the detailed interrogation of lineage and mutational rate differences between these tissues^17,18^.

Finally, to determine if this approach could be applied to other forms of genetic variation, we tested germline mosaicism of two structural *de novo* variant classes: 1) large deletions and duplications, and 2) short tandem repeat (STR) expansions and contractions. F21 and F22 both harbor likely pathogenic *de novo* structural variants: a ~1.5 Mb duplication (F21) and a 130 kb deletion (F22) (Fig. 4a)^12^. While the duplication in F21 and a non-pathogenic, additional deletion in F22 did not appear to be mosaic in sperm (Extended Data Fig. 8d-g), we found evidence of sperm-specific mosaicism for the pathogenic deletion in F22 using read depth and split-read information (Extended Data Fig. 8a,b). This was confirmed with a PCR based strategy, and copy number detection by ddPCR estimated 0.15 deletion copies in paternal sperm, or an effective AF of ~7.5% given the presence of an unaffected reference allele in the genome (Fig. 4b-d and Extended Data Fig. 8c).

**Figure 4.**
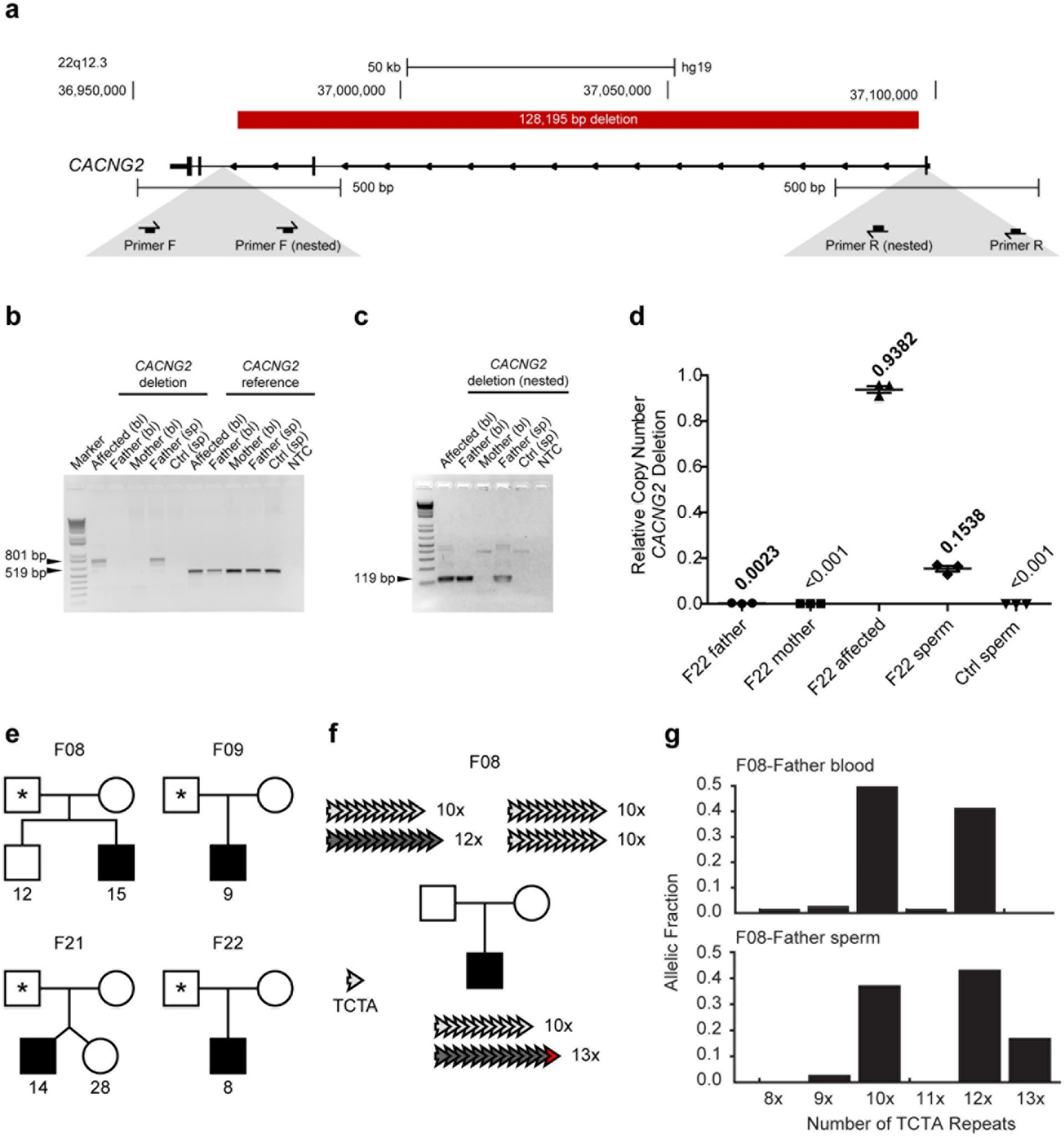
Germline mosaicism extends to structural variants. **a**, Schematic depicting part of the genomic locus of *CACNG2* and the pathogenic 128,195 bp deletion found in family F22. Below: primers used for the nested PCR to detect the deletion. **b**, Agarose gel resolving theprimary PCR products from the indicated individuals from blood (bl) and sperm (sp). *CACNG2* deletion: PCR spanning the deletion locus to amplify an 801 bp band if deletion is present. *CACNG2* reference: PCR within the deleted locus to amplify a 519 bp band. The deletion-specific band was detected in the child’s blood and the father’s sperm sample. **c**, Agarose gel resolving the nested PCR products arranged as in **b**. Note that this strategy also showed positive signal for the paternal blood. Together with **b**, this suggested that the deletion allele is present at low AF in the paternal blood and at considerably higher levels in the paternal sperm. **d**, Copy number quantification of the *CACNG2* deletion by ddPCR. Samples from father, mother, and child were derived from blood (and sperm in the case of the father). The deletion allele was present at a copy number of 0.1538 in paternal sperm, which consequently means that it was present at an AF of ~7.5%. **e**, Schematic showing all four pedigrees for F08, F09, F21, and F22 and the number of detected *de novo* short tandem repeat variants (dSTRΔs) for each child. **f**, Schematic showing an example of a dSTRΔ in F08, where the child had an expansion of a tetranucleotide repeat (TCTA) on the paternal haplotype (12x to 13x). **g**, Detailed analysis of the TCTA repeat numbers in paternal blood and sperm reveals a sperm-specific mosaicism of the 13x repeat at an AF of ~17.5%.

Short tandem repeats (STRs) are particularly dynamic in the genome and can expand or contract *de novo* during transcription, replication, and meiosis, impacting gene functions and causing diseases such as Huntington’s disease or Fragile X syndrome^19,20^. As *de novo* short tandem repeat changes (dSTRΔs) have to date not been implicated in ASD, we could not identify likely pathogenic variants. Therefore, we adopted a strategy similar to the one used to evaluate dSNVs: calling non-pathogenic dSTRΔs using WGS data and then evaluating their presence in paternal sperm and blood (Fig. 2a and 4e). We detected 86 non-pathogenic dSTRΔs, five of which were exclusively mosaic in the paternal sperm at an AF ranging from 17.5% to 1.9% (Fig. 4e, Extended Data Fig. 9a-e, and Supplementary Table 5). The dSTRΔ with the highest AF was a tetranucleotide repeat expansion in the child (Fig. 4f). It was not present in the somatic tissues of either parent, but was detected in the father’s germline with a 17.5% AF (Fig. 4g and Extended Data Fig. 9f-g). Highly unstable STR pre-mutations may be prone to early mosaicism. Assessing dSTRΔs in the germline in pre-mutation carriers may help to adjust the individualized risk to offspring.

This study is the first, to our knowledge, to directly assess this type of mosaicism in a relevant germline tissue, paternal sperm. While previous reports have estimated germline mosaicism from dSNV recurrence among siblings or detection in peripheral tissues^9,10,21,22^, here we directly measure germline mosaicism in males and present a conceptual framework for refining disease risk to offspring. We show that more than 3% of dSNVs are mosaic in the paternal germline. Although we focus on families with ASD, these findings are applicable to a range of diseases caused by *de novo* mutations. The analysis of sperm instead of non-germline tissues could be an important addition to clinical practice allowing for accurate prediction of recurrence risk, but also for detection of previously non-transmitted, mosaic mutations prior to conception.

## Methods

### Patient recruitment

Patients were enrolled according to approved human subjects protocols at the University of California for blood, saliva, and semen sampling. Semen was collected for all fathers of families F01-12 and F21-22. For F01-04 we obtained saliva from the fathers and their family members, for F05-12 and F21-22 we extracted DNA from blood. WES trio analysis for F01-F04 was performed on DNA extracted from lymphocyte cell lines (generated by the NIMH Repository) and results were confirmed in saliva samples, WGS trio analysis for F04-12 and F21-22 was performed on DNA derived from blood.

### WES and WGS trio analysis

Exome capture and sequencing of F01-F04 was performed at the New York Genome Center (Agilent Human All Exon 50 Mb kit, Illumina HiSeq 2000, paired-end: 2x100) and the Broad Institute (Agilent Sure-Select Human All Exon v2.0, 44Mb baited target, Illumina HiSeq 2000, paired-end:2x76). Sequencing reads were aligned to hg19 reference using BWA (v0.7.8)^23^. Duplicates were marked using Picard’s MarkDuplicates (v1.83, http://broadinstitute.github.io/picard) and reads were re-aligned around INDELs with GATK’s IndelRealigner^24^. Variant calling for SNVs and INDELs was according to GATK’s best practices by first calling variants in each sample with HaplotypeCaller and jointly genotyping them across the entire cohort using CombineGVCFs and GenotypeGVCFs. Variants were annotated with SnpEff (v4.2)^25^ and SnpSift (v4.2)^26^ and allele frequencies from the 1000 Genomes Project and the Exome Aggregation Consortium (ExAC)^27,28^. *De novo* variants were called for probands using Triodenovo^29^ with a minimum *de novo* quality score (minDQ) of 2.0 and subjected to manual inspection. WGS sequencing and analysis for F05-F12 and F21-22 were performed as described previously^3,12^.

### Blood and saliva extraction

DNA was extracted on an Autopure LS instrument (Qiagen, Valencia, CA).

### Sperm extraction

Extraction of sperm cell DNA from fresh ejaculates was performed as previously described^30^. In short, sperm cells were isolated by centrifugation of the fresh ejaculate over an isotonic solution (90%) (Sage/Origio, ART-2100; Sage/Origio, ART-1006) using up to 2 mL of the sample. Following a washing step, quantity and quality were assessed using a cell counting chamber (Sigma-Aldrich, BR717805-1EA). Cells were pelleted and lysis was performed by addition of RLT lysis buffer (Qiagen, 79216), Bond-Breaker TCEP solution (Pierce, 77720), and 0.2 mm stainless steel beads (Next Advance, SSB02) on a Disruptor Genie (Scientific Industries, SI-238I). The lysate was processed using reagents and columns from an AllPrep DNA/RNA Mini Kit (Qiagen, 80204). Concentration of the final eluate was assessed employing standard methods. Concentrations ranged from ~0.5-300 ng/μl. Sperm extracted DNA was stored on −20°C.

### Sanger sequencing of SNVs

PCR and Sanger sequencing were performed according to standard methods. Primer sequences can be found in Supplementary Table 6. Validated mutations and surrounding SNPs were also used as basis for the design of ddPCR assays where applicable.

### ddPCR design, validation, and setup of experiments

Using the Primer3Plus web interface^31–33^, the amplicon and probes for wild-type and mutant were designed to distinguish reference and alternate allele (settings in Supplementary Document 1). Probes were required to be located within 15bp up- and 15 bp downstream of the mutation and adjusted, so melting temperatures (Tm) were matched between reference and alternate probe. In addition, if possible, amplicons were kept at 100 bp or shorter and probes at 20 bp or shorter. Specificity of the primers was assessed using Primer-BLAST^34^. Custom primer and probe mixes (primer to probe ratio of 3.6) were ordered from IDT with FAM-labeled probes for the alternate, and HEX-labeled probes for the reference allele (Supplementary Table 6). Optimal annealing temperature, specificity, and efficiency were tested using custom gblocks (IDT) or patient DNA at a range of dilutions. ddPCR was performed on a BioRad platform, using a QX200 droplet generator, a C1000 touch cycler, a PX1 PCR Plate Sealer, and a QX200 droplet reader with the following reagents: ddPCR Supermix (BioRad, 1863024), droplet generation oil (BioRad, 1863005), cartridge (BioRad, 1864008), and PCR plates (Eppendorf, 951020346). Aiming for 30-60 ng per reaction, up to 8 μl of DNA solution were used in a single reaction. Data analysis was performed using the software packages QuantaSoft and QuantaSoft Analysis Pro (BioRad). Each run included technical triplicates. For direct comparison of sperm samples we used seven technical replicates, except for F01, where the total amount of sperm DNA was limiting. Across all ddPCR reactions that were designed for SNV detection, we determined that the minimum AF that we could reliably detect was 0.1%. Therefore, we set this as threshold of detection. Raw data for ddPCR experiments can be found in Supplementary Table 7.

### Data processing

Graphs were generated and data analyzed using GraphPad Prism, R, and Python (matplotlib library).

### WGS of matched sperm and blood samples

WGS was performed using an Illumina TrueSeq PCR-free kit (350bp insertion) on an Illumina HiSeqX. Paired-end FASTQ files of deeply (>90x) sequenced blood and sperm samples from fathers were aligned to the hg19 reference genome (1000Genomes version 37) with bwa mem (version 0.7.15-r1140), specifying the–M option that tags chimeric reads as secondary, required for some downstream applications that implement this legacy option. The resulting average coverage was 117x for blood samples and 109x for sperm samples with an average read length of 150bp for both sets. Duplicates were removed with the markdup command from sambamba (version 0.6.6), and base quality scores were recalibrated with the Genome Analysis ToolKit (GATK version 3.5-0-g36282e4). SNPs and INDELs were called with HaplotypeCaller jointly genotyping within pedigrees, consisting of the deep coverage (>90x) genomes from father’s blood and sperm and 40x coverage genomes derived from blood of the parents, sibling (F08 and F21 only), and proband.

### Oxford Nanopore sequencing and analysis

We generated whole genome sequencing libraries with Oxford Nanopore 1D long reads for four affected probands (Families: F08, F09, F21, and F22) according to manufacturer’s recommendations. FASTQs were aligned to the hg19 reference genome with bwa mem with the ‘-x; ont2d’ option for ONP reads. Coverage of proband samples ranged from 15x; to 3x (average 9x) with average read length ranging from 7,839bp to 4,645bp (average: 6,777bp).

### Haplotype phasing

To determine dSNV phase, we first identified a set of phase-informative SNPs using the germline variant calls from our 40x WGS data. Phase-informative SNPs were those where the child was heterozygous and either 1) one parent was heterozygous or homozygous for the alternate allele while the other parent was homozygous for the reference allele, or 2) one parent was heterozygous while the other parent was homozygous for the alternate allele. Second, we identified long-reads (Oxford Nanopore reads, average length 6,777 bp) that contained both a dSNV and one or more phase-informative SNPs. We then counted the number of dSNV and phase-informative SNP combinations that were present in reads and consistent with the dSNV occurring on a maternal or paternal haplotype. Reads containing an INDEL flanking either the dSNV or the phase-informative SNP were excluded from the analysis. Finally, we assigned the dSNVs to maternal and paternal haplotypes if there were: 1) a minimum of two counts, and 2) the haplotype with the majority of counts had at least 2/3 of total counts. Out of the 256 variants from the four affected children, we succeeded in phasing for 187 (73.0%), of which 152 were phased to the paternal haplotype (81.3%; α~4). These paternal dSNVs were then used for further analysis as described below.

### Mosaic dSNV analysis

Using the read information generated by HaplotypeCaller, we determined AF for previously called dSNVs. We additionally annotated dSNVs that fell in repetitive regions of the human genome using the repeatMasker (rmsk.txt) file from UCSC. We manually filtered those variants that were homozygous in the reference and heterozygous in the proband, as well as variants that were present in both blood and sperm at AF that suggested an inherited heterozygous SNP (i.e. AF>35% in both blood and sperm). This resulted in a total of 355 dSNVs that were analyzed, 152 being paternally phased (see phasing methods). Out of 355 variants, 169 were outside of repetitive regions. Separate analysis of these, revealed similar rates of mosaicism (Supplementary Table 3). Thus, we concluded that assessment of all variants is acceptable for this approach. Out of the total of 355, 59 (all) and 12 (AF>2%) were showing read evidence in sperm, blood, or both. 7 (all) and 2 (AF>2%) of these were phased to the maternal haplotype (i.e. were most likely false positives). Out of the paternally phased 152 variants, 29 (all) and 9 (AF>2%) were showing read evidence in sperm, blood, or both. Mosaic variants were categorized based on their presence or absence in sperm and blood. To be called sperm enriched, a variant’s AF had to be three times higher in sperm than in blood (α>3).

### MuTect/Strelka mosaic variant calling

Sequencing reads for four pairs of blood and sperm samples were aligned to the hg19 version of the reference genome using the iSAAC aligner^35^ using the option --base_quality_cutoff 15. Duplicates were marked with Picard’s MarkDuplicates (v1.128, http://broadinstitute.github.io/picard) and INDELs were realigned using GATK’s IndelRealigner (v3.5)^24^. We then called sperm- and blood-specific SNVs using two somatic variant callers with default parameters, Strelka (v2.7.0)^36^ and muTect (v3.1)^37^, setting the sperm sample as “tumor” and the blood sample as “normal”. For blood specific-variants, we did the reverse. We defined a high threshold for somatic calls for each sperm-blood and blood-sperm comparison by taking the intersection of variants identified by both Strelka and MuTect. These high quality calls were further filtered to reduce potential false positives as follows. We removed calls that fell into repetitive regions, using the RepeatMasker (rmsk.txt) file from UCSC, and removed calls that fell within 5 bp of a germline INDEL. For mosaic variant analysis in blood, F09 was an outlier with respect to number of variants that were called. Consequently, analyses were performed with and without variants from this individual to reflect this issue.

### Mosaic SV analysis of WGS data

We searched for evidence for mosaicism in the fathers using depth of coverage, split-reads, discordant paired-ends, and B-allele frequency in deeply sequenced paired-end genomes. Depth of coverage was estimated as the median per base-pair coverage within the SV locus, while omitting positions that overlapped assembly gaps, RepeatMasker elements, short tandem repeats, and segmental duplications. We estimated copy number by dividing the median depth of coverage by the median coverage of the chromosome and multiplying by 2. Split-reads (also known as chimeric reads) are those with multiple alignments to the genome. If a read spanned a deletion or tandem duplication breakpoint, two alignments were generated with each segment mapping to opposite ends of the breakpoint. Similar to split-reads, discordant paired-ends had read fragments that span the SV breakpoint, but the SV breakpoint resided in the unsequenced insert of the fragment. Consequently, the paired-ends mapped to opposite ends of the breakpoint producing an insert size approaching the size of the SV. We searched ±250 bp from the predicted breakpoint for SV supporting reads, which were unique reads that were either split or contained discordant paired-ends with breakpoints that overlap at least 95% reciprocally to the SV. We reported the proportion of supporting reads to non-informative reads (those that do not support the SV) within the +/-250bp windows, which roughly estimates proportion of mosaicism. Additionally for the *de novo* duplication SV, we searched for deviations in B-allele frequency defined as the proportion of reads that support the alternate variant to all reads covering the variant in question.

### Mosaic SV analysis using PCR and ddPCR

Nested PCR was performed using blood DNA extracted from the F22 trio (proband, mother, and father), as well as sperm from the F22 father and a non-related male. Primers were designed using Primer3Plus online software^31^ to span the deletion breakpoints within *CACNG2* determined by WGS analysis within 500 bp windows up- and down-stream of the predicted deletion. Additionally, a reverse primer was designed to be used with the nested forward primer as an amplification control (Supplementary Table 6). All PCR reactions were 25 μl volumes and included 20 mM Tris-HCl (pH 8.4), 50 mM KCl, 2 mM MgCl2, 1 U of Taq (Thermo Fisher Scientific, Waltham, MA), and 300 nM of each appropriate primer. DNA template was 50 ng of DNA from blood or sperm for the initial PCR (using the external set of primers), or 1 μl of the initial PCR product for the nested (internal) PCR. PCR reactions were run following a standard ramp speed protocol using a C1000 Touch™ Thermal Cycler (Bio Rad, Hercules, CA) with cycling consisting of a 2 min initiation at 95°C, 35 cycles of 95°C for 30 s, 55°C anneal for 30 s, and 72°C for 1 min, followed by a final extension at 72°C for 3 min. Products were resolved on 2% agarose gels. For ddPCR analysis, primer and probe sets for F22 were designed using Primer3Plus (Supplementary Document 1 and Supplementary Table 6). Probe annealing temperature was designed to be 5°C higher than the primer binding temperatures. Primers were designed to span the deletion breakpoints within *CACNG2.* A custom primer and FAM-labeled probe mix at a primer:probe ratio of 750 nM:250 nM was ordered from Bio Rad (Hercules, CA) as well as a HEX-labeled pre-validated copy number variation assay specific for *RPP30* as an internal control (assay ID: dHsaCP2500350). ddPCR was performed and analyzed as described above. Raw data for ddPCR experiments can be found in Supplementary Table 8.

### dSTRΔ calling and mosaicism detection

For the analysis of STR expansions and contractions, we used HipSTR^38^ (version v0.2-311-g9bcd580) jointly on all BAM files (40x trios and >90x blood and sperm of fathers). We used the reference STR set provided by HipSTR for GRCh37 (GRCh37.hipstr_reference.bed) and default options except for: --def-stutter-model and --output-gls. We further ran HipSTR’s denovofinder tool on each of the 40x trios with the option --uniform-prior. The following, strict filters were applied for the detection of a *de novo*: required genotype call in all family members; posterior probability of de novo mutation ≥0.9; ignored mutations that are not a multiple of the repeat unit; ignored if allele lengths followed Mendelian inheritance or if *de novo* allele also was found in one of the parents; minimum genotype quality of 0.9 in all family members; minimum percentage of reads with stutter or INDEL was 20% for all family members; required at least 10 spanning reads in all family members; required at least 20% of reads to support each allele in each family member; new allele was excluded if homozygous in the child; removed segmental duplications (UCSC segmental duplication track)^39,40^; and removed calls that overlapped with >10 entries in DGV^41^. We then annotated the remaining loci with their frequencies in the >90x sperm and blood samples. We calculated the posterior probability of a *de novo* mutation using HipSTR outputs of no mutation, *de novo* mutation, and other. We converted this to a posterior assuming the following priors: prob(mutation)=0.0001 and prob(other)=0.01. dSTRΔs were qualified as inconclusive if mosaicism was detected in mother and father or only in paternal blood; as true *de novo* if no mosaicism was detected in the parents; as maternal if mosaicism was only detected in the mother; and as paternal if mosaicism was detected in blood and sperm, or sperm only.

## Acknowledgements

We thank the participants in this study for their contribution. This study was supported by NIH U01MH108898, R01NS083823, the Simons Foundation Autism Research Initiative (SFARI) and the Howard Hughes Medical Institute (to J.G.G.). Sequencing support was provided by the Rady Children’s Institute for Genomic Medicine and Oxford Nanopore. O.D. acknowledges support from the Silverman Family Foundation and Finding A Cure for Epilepsy and Seizures. M.W.B. is supported by an EMBO Long-Term Fellowship (ALTF 174-2015), which is co-funded by the Marie Curie Actions of the European Commission (LTFCOFUND2013, GA-2013-609409).

## Author Contributions

M.W.B, J.G.G., and J.S. conceived the project and planned the experiments. M.W.B., M.K., L.L.B., C.A., and A.N. performed the experiments. R.D.G. and D.A. performed the bioinformatic analysis. D.M., R.K., and E.S. performed the *de novo* analysis of the cohort collected and provided by O.D. K.N.J., O.H., J.M.-V., and M.W.B. requested, organized, and handled patient samples. M.W.B., J.G.G., and J.S. wrote the manuscript with input from R.D.G. and K.N.J. All authors have seen and commented on the manuscript prior to submission.

## Author Information

M.W.B., K.N.J., J.S., and J.G.G. are inventors on a provisional patent (62/512,368) filed by UC, San Diego that covers the work in this manuscript. Correspondence and requests for materials should be addressed to

---

**Extended Data Figure 1.**
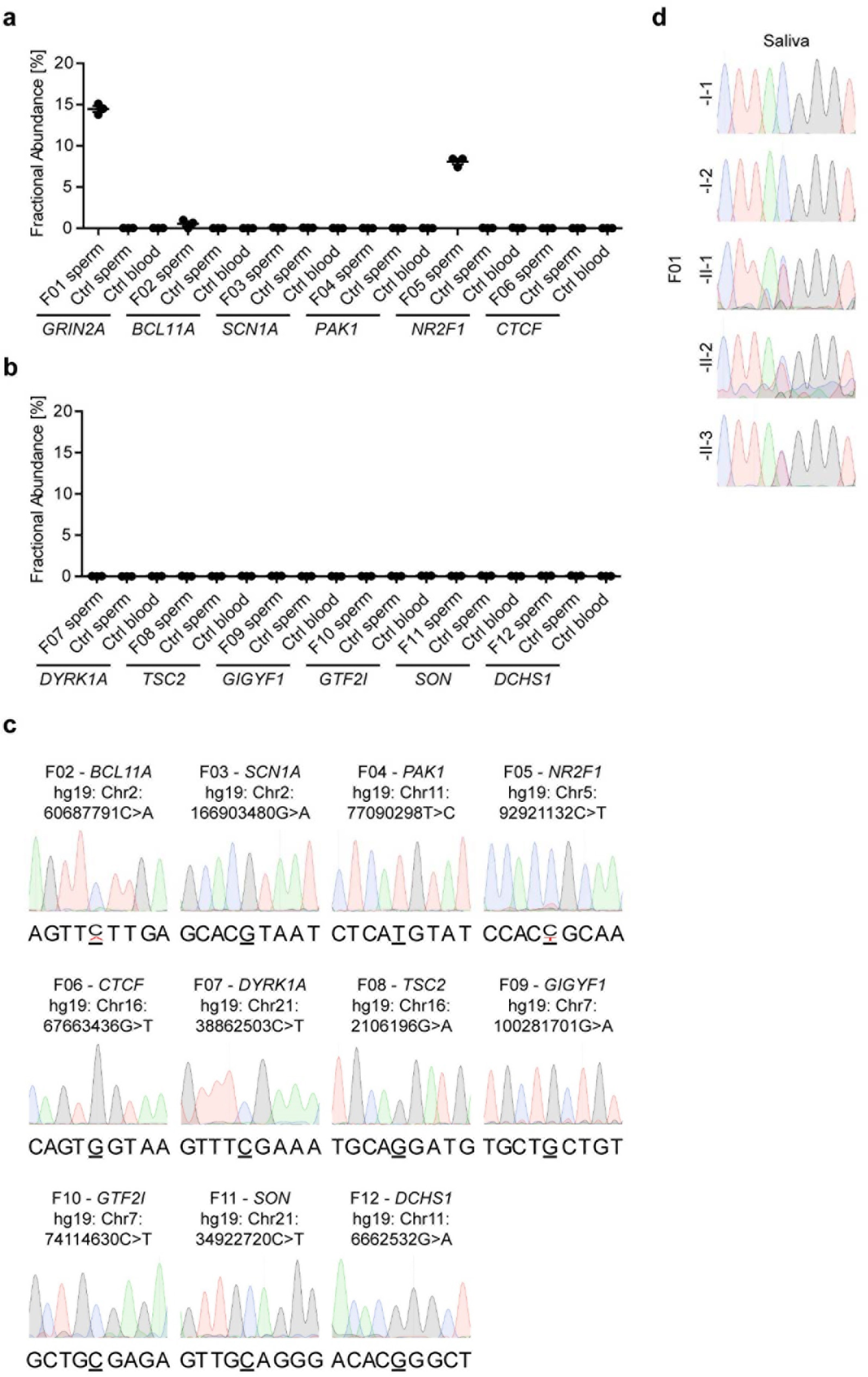
Detection of paternal germline mosaicism in 3 out of 12 ASD families. **a**,**b**, Fractional abundance (determined by ddPCR) of the mutant allele in paternal sperm for the relevant dSNV in the 12 families. Ctrl-an unrelated sperm or blood sample, as indicated, acting as control. Graphs show individual data points (technical triplicates) and mean ± SEM. **c**, Sanger sequencing results showing the locus harboring the dSNV for each family. Confirming the ddPCR results, F02 and F05 showed mosaicism at their respective positions. **d**, Sanger sequencing results showing the C>T conversion locus in *GRIN2A* in F01 for all family members. The mutation was absent in the saliva of both parents, but present as a heterozygous allele in all 3 children. Parts of this panel are shown in Fig. 1e.

**Extended Data Figure 2.**
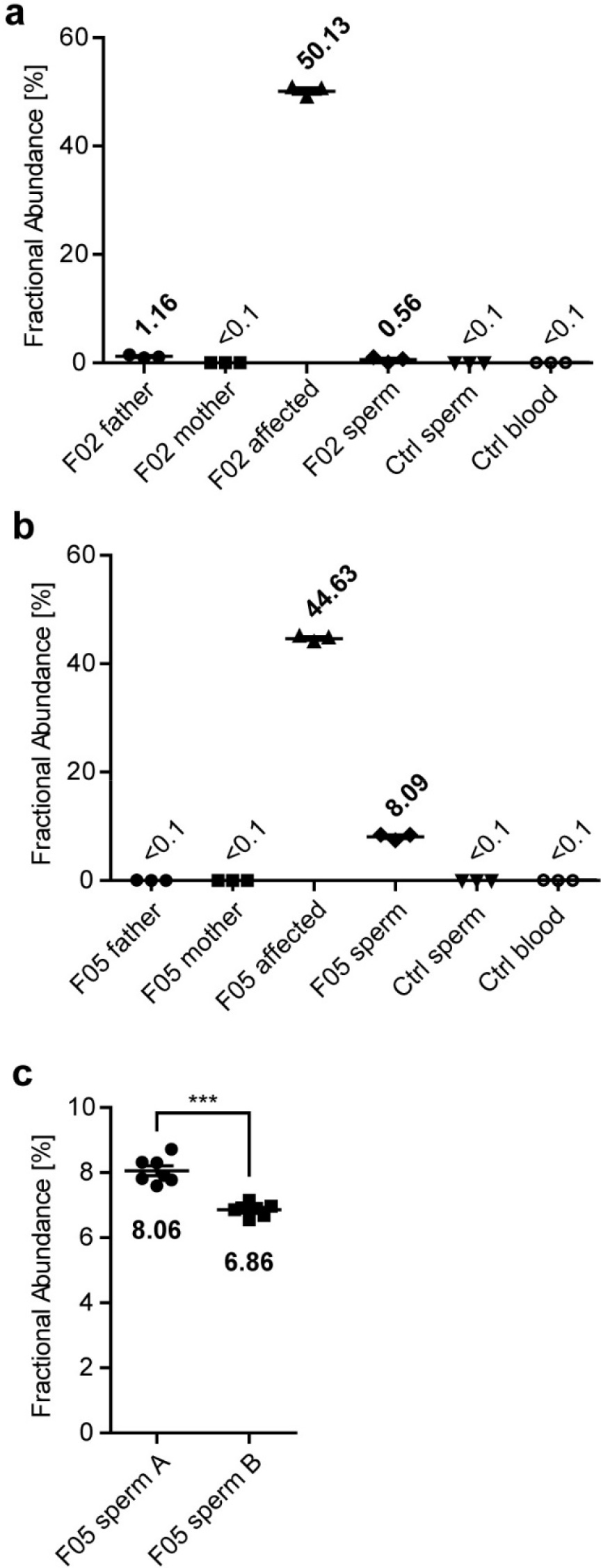
Extended ddPCR results for the mosaic variants in F02 and F05. **a**, Fractional abundance (determined by ddPCR) of the mutant *BCL11A* allele in F02. While the variant was absent from the saliva sample from mother, as well as the blood and sperm control (ctrl) samples, it was present at low levels in the father’s saliva and sperm. This variant was detected as mosaic at similar levels in both tissues, although the very low sperm yield obtained may have influenced the results. **b**, as in **a**, but for F05. Mosaicism could only be detected in the paternal sperm, but not the blood sample. **c**, Fractional abundance (determined by ddPCR) comparing two biological replicates of paternal sperm from F05. Sperm sample A (used for ddPCR analysis in Fig. 1, Extended Data Fig. 1, and Extended Data Fig. 2b) was significantly different from sample B, suggesting variation of mosaicism over time. ***P<0.001 (unpaired t-test, two-tailed). Graphs in **a-c** show individual data points (technical triplicates in **a**,**b** and seven technical replicates in **c**) and mean ± SEM.

**Extended Data Figure 3.**
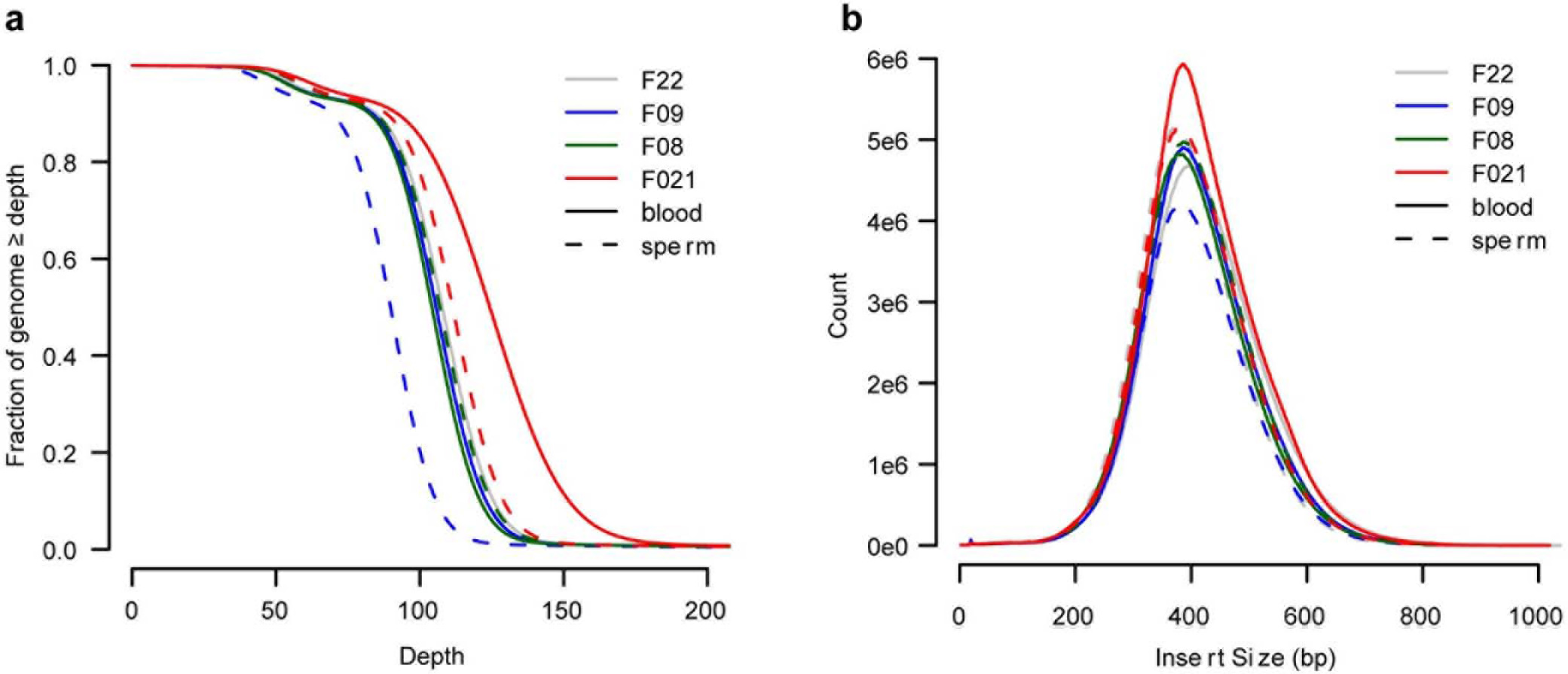
Whole genome sequencing metrics. **a**, Plot showing the read depth for the blood and sperm samples from the fathers of F08, F09, F21, and F22. **b**, Plot showing the insert size distribution for the same data sets as in **a**.

**Extended Data Figure 4.**
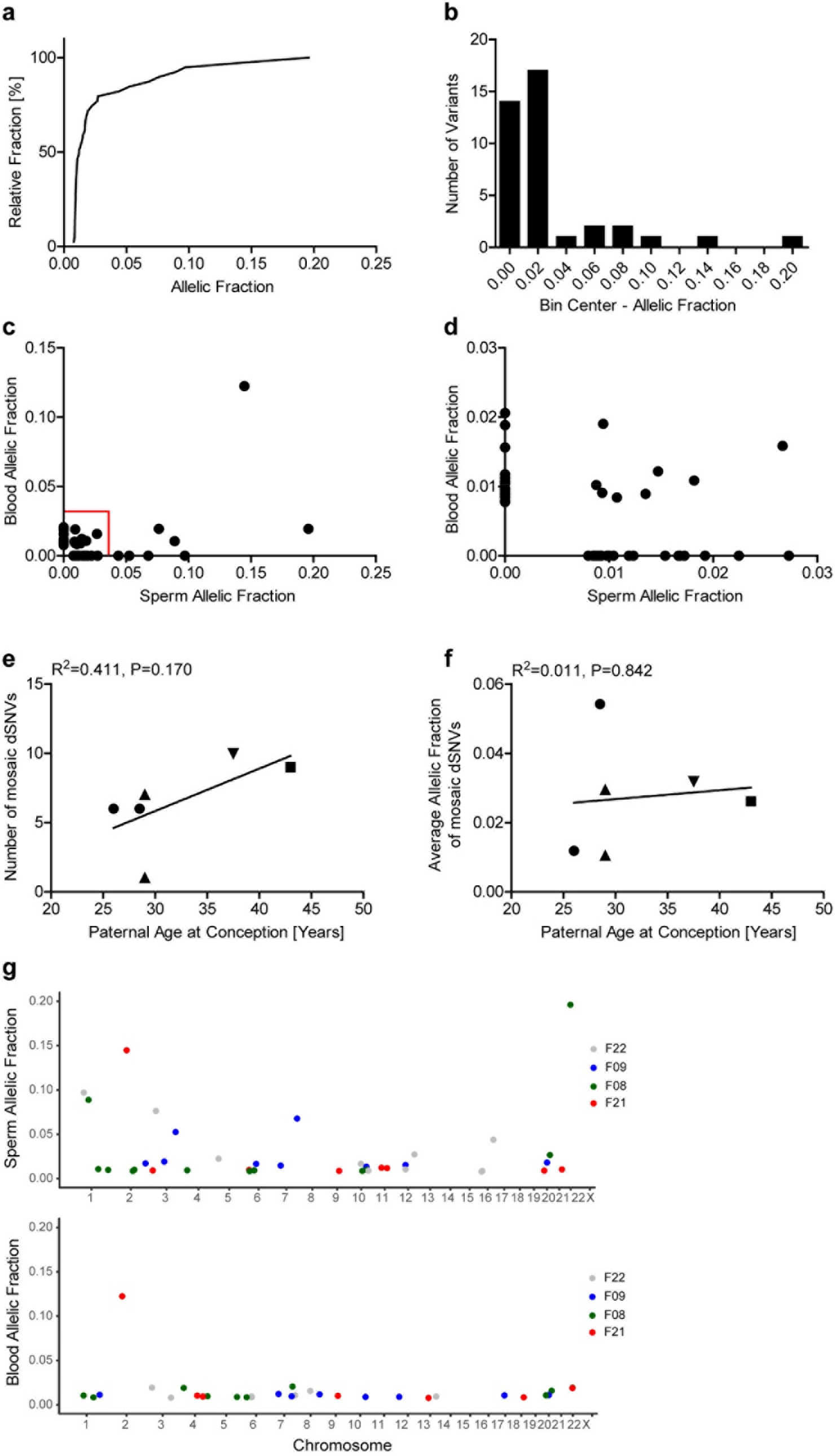
Supplementary graphs for the mosaicism analysis of dSNVs. **a**, Plot showing the cumulative relative fraction of mosaic dSNVs in sperm. **b**, Frequency distribution plot for same data as in **a**. **c,d**, Plot showing the AF in sperm and blood for all mosaic dSNVs. **d** is a magnification of the red box in **c**. **e**, Plot showing the number of mosaic dSNVs present at the paternal age at conception. Line shows a regression curve suggesting a positive correlation that is non-significant (R^2^=0.411, P=0.170). **f**, Plot showing the average AF of mosaic dSNVs relative to the paternal age at conception. Line shows a regression curve without positive correlation (R^2^=0.011, P=0.842). **g**, Plot of all mosaic variants denoting their positions on the chromosomes for sperm and blood.

**Extended Data Figure 5.**
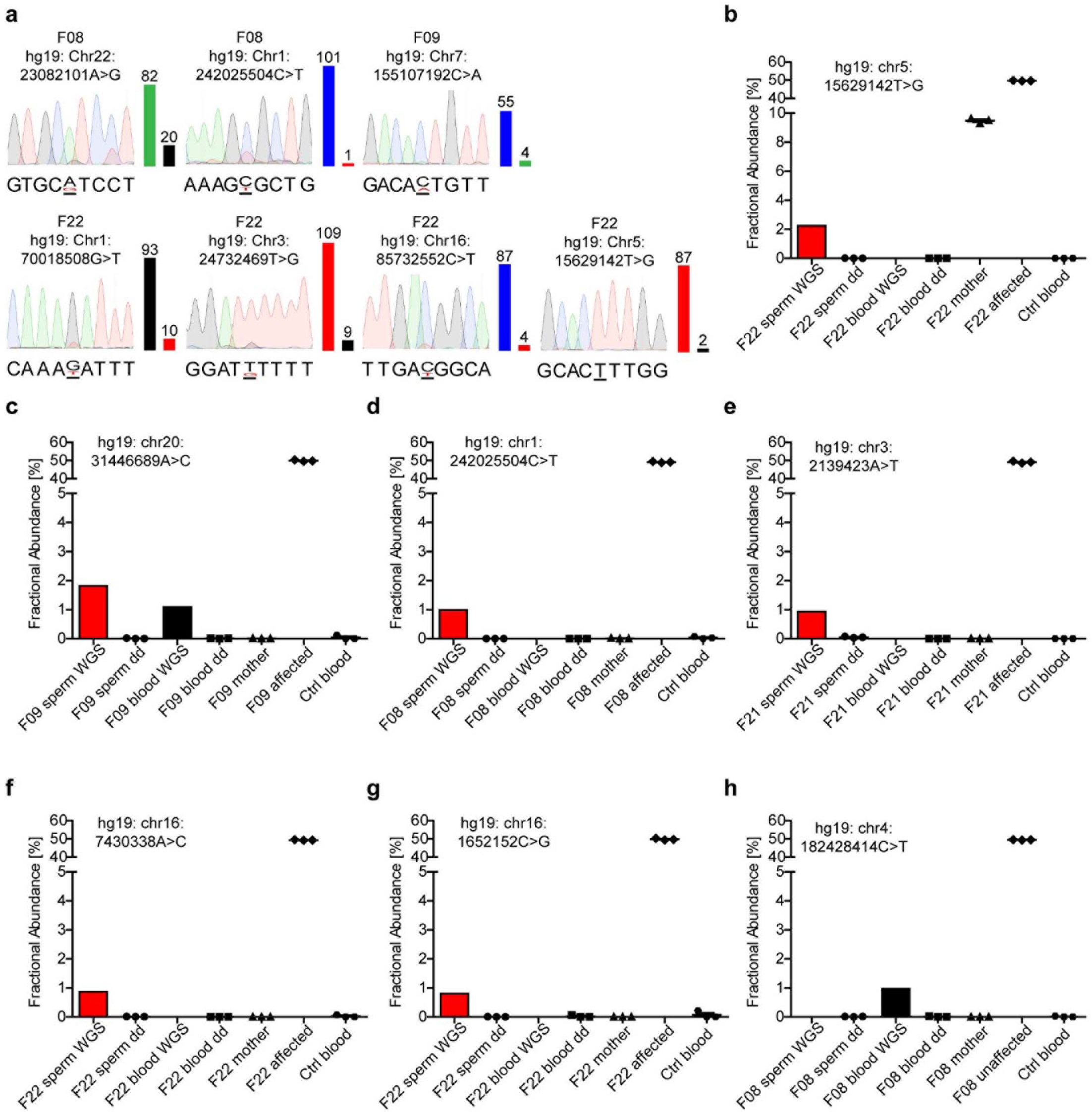
Orthogonal confirmation experiments of mosaic dSNVs. **a**, Sanger sequencing results for seven of the detected mosaic variants, showing mosaicism in six cases. Numbers and bars beside the chromatograms depict read counts in the WGS data for the reference allele on the left and the mutant allele on the right. Variant F08:hg19: Chr1:242025504C>T showed suspiciously high mosaic levels in the chromatogram compared to the WGS results. **b-h**, Fractional abundance (determined by ddPCR or WGS read counts) of the indicated mutant alleles. Mother and (un)affected indicate blood samples from the mother and the child that harbored this mutation. Note that none of the low mosaic variants showed mosaicism in paternal sperm. Variant F22: chr5:15629142T>G depicted in **b** exhibited mosaicism at ~10% in the mother, consistent with its phasing to the maternal haplotype (Supplementary Table 3). Variant F08:hg19: Chr1:242025504C>T (depicted in **d**) was also interrogated in **a** and showed the surprisingly high levels of mosaicism. These data suggested that the Sanger sequencing result was a false positive, probably caused by repetitive sequences. Taken together, orthogonal quantification by ddPCR suggested that low level variants are highly unreliable as we could not confirm them with orthogonal methods. Graph shows individual data points (technical triplicates) and mean ± SEM for the ddPCR data.

**Extended Data Figure 6.**
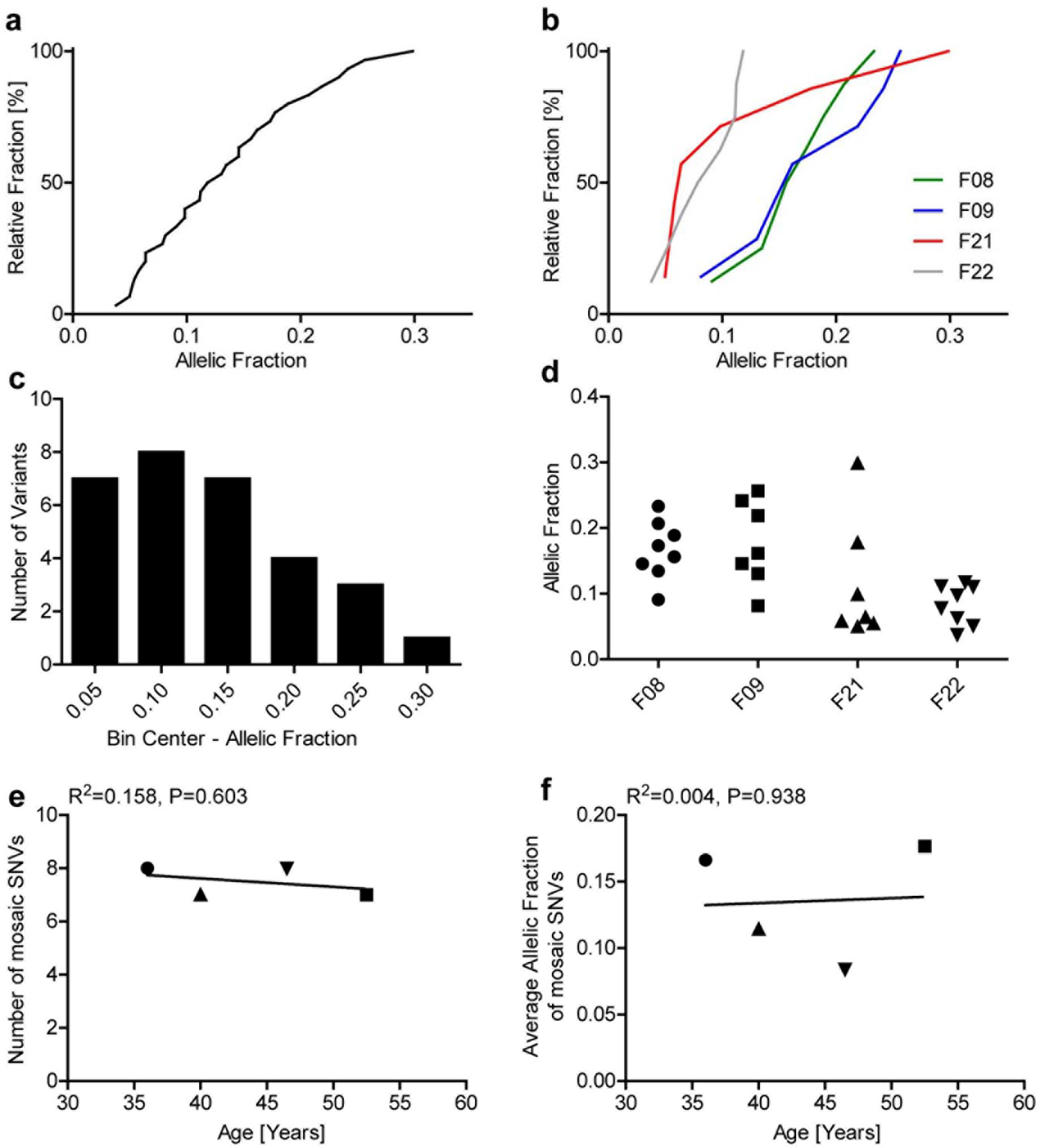
Supplementary graphs for the mosaicism analysis of SNVs by MuTect and Strelka. **a**, Plot showing the cumulative relative fraction of mosaic SNVs detected in sperm employing our MuTect/Strelka pipeline. **b**, As in **a**, but separated by origin of the variants. **c**, Frequency distribution plot for same data as in **a**. **d**, As in **b**, but showing the AF distribution per sample. **e**, Plot showing the number of mosaic SNVs relative to the paternal age at sample collection. Line shows a regression curve without correlation (R^2^=0.158, P=0.603).**f**, Plot showing the average AF of mosaic SNVs relative to the paternal age at sample collection. Line shows a regression curve without correlation (R^2^=0.004, P=0.938).

**Extended Data Figure 7.**
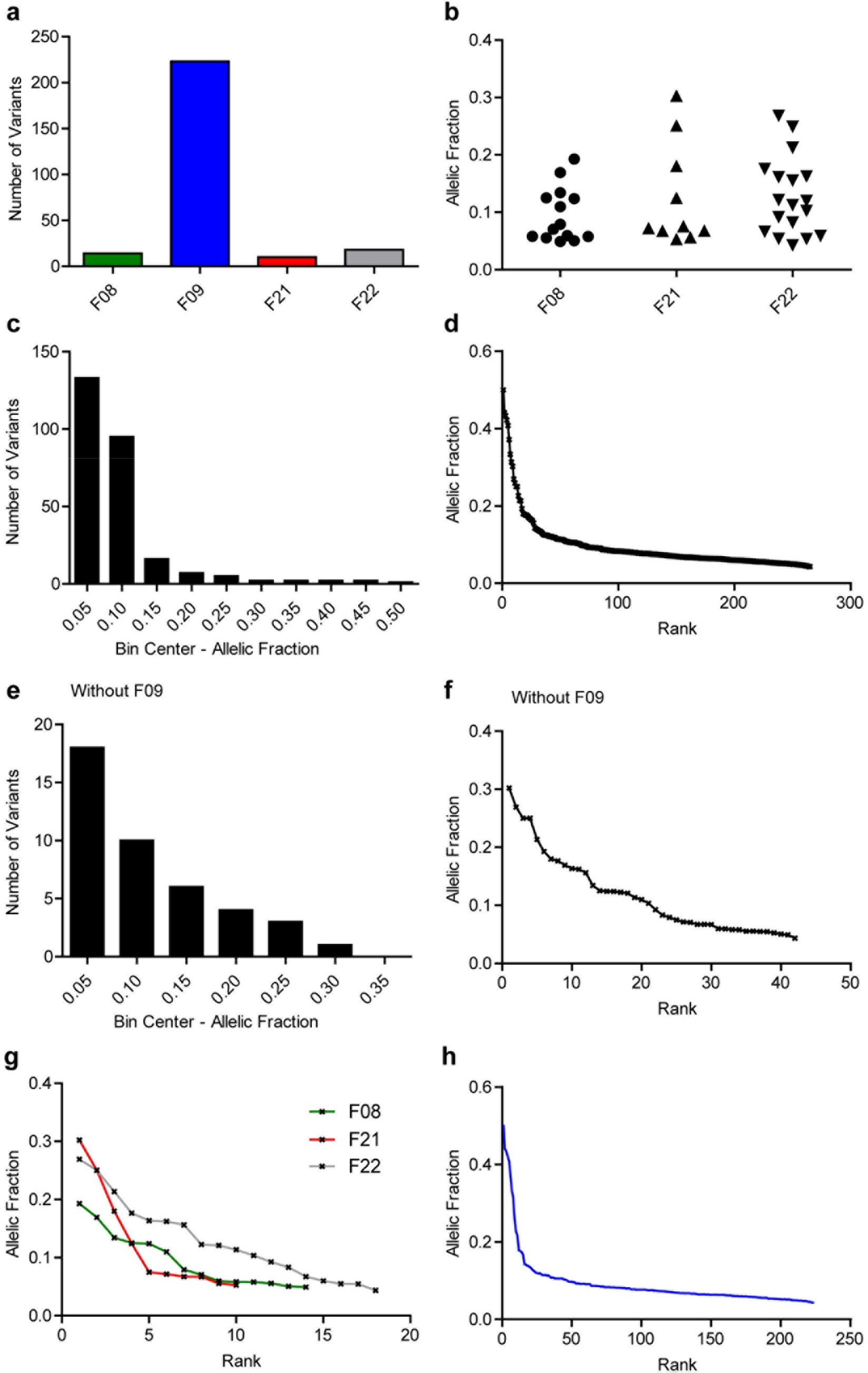
Unbiased analysis of mosaic variants in blood. **a**, Plot showing the number of variants detected per sample. F09 showed an aberrantly high number of variants relative to the other individuals. **b**, Plot showing the AF distribution per sample (without F09). **c**, Frequency distribution of all mosaic SNVs found in blood. **d**, Plot showing all mosaic SNVs found in blood and their AF. **e,f**, Same as **c,d**, but without F09. **g,h**, Plot showing the mosaic SNVs found in blood and their AF by origin for F08, F21, and F22 (**g**), as well as F09 (**h**).

**Extended Data Figure 8.**
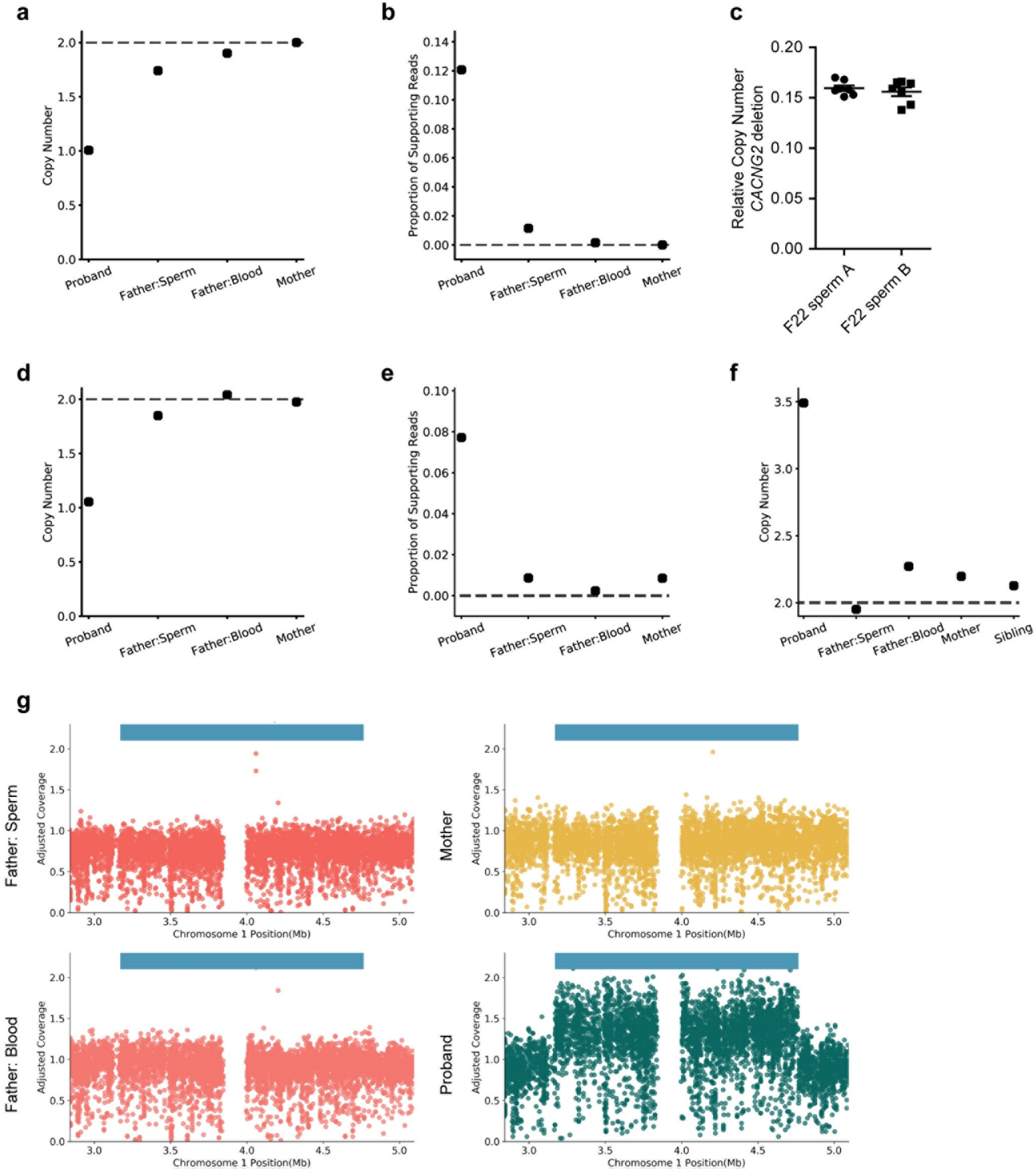
Structural variant detection in the WGS data sets for F21 and F22. **a**, Copy number analysis of the region deleted in the structural variant (SV) depicted in Fig. 4a. Both the father’s sperm and blood showed a reduction relative to the mother. **b**, Supporting split read data for the same deletion further supported mosaicism in sperm. **c**, Copy number quantification of the *CACNG2* deletion by ddPCR. Two biological replicate sperm samples from the father of F22 were compared to each other. Sperm sample A was the one used for the analysis shown in Fig. 4b-d and >90x WGS. There was no significant difference between the two samples (unpaired t-test, two-tailed). **d,e**, same data as in **a** and **b** for a separate, non-pathogenic deletion found in F22. Although copy number did support mosaicism in the paternal sperm, split read evidence did not, as all the positive reads in the paternal sperm and the mother were faulty alignments (data not shown). Together, these data suggest that this deletion is not mosaic in sperm. **f,g**, Copy number analysis of a 1.6 Mb duplication on chromosome 1 in F21 that is likely pathogenic. Overall copy number (**f**) and detailed locus analysis (**g**) both suggest that this variant is not mosaic in paternal sperm.

**Extended Data Figure 9.**
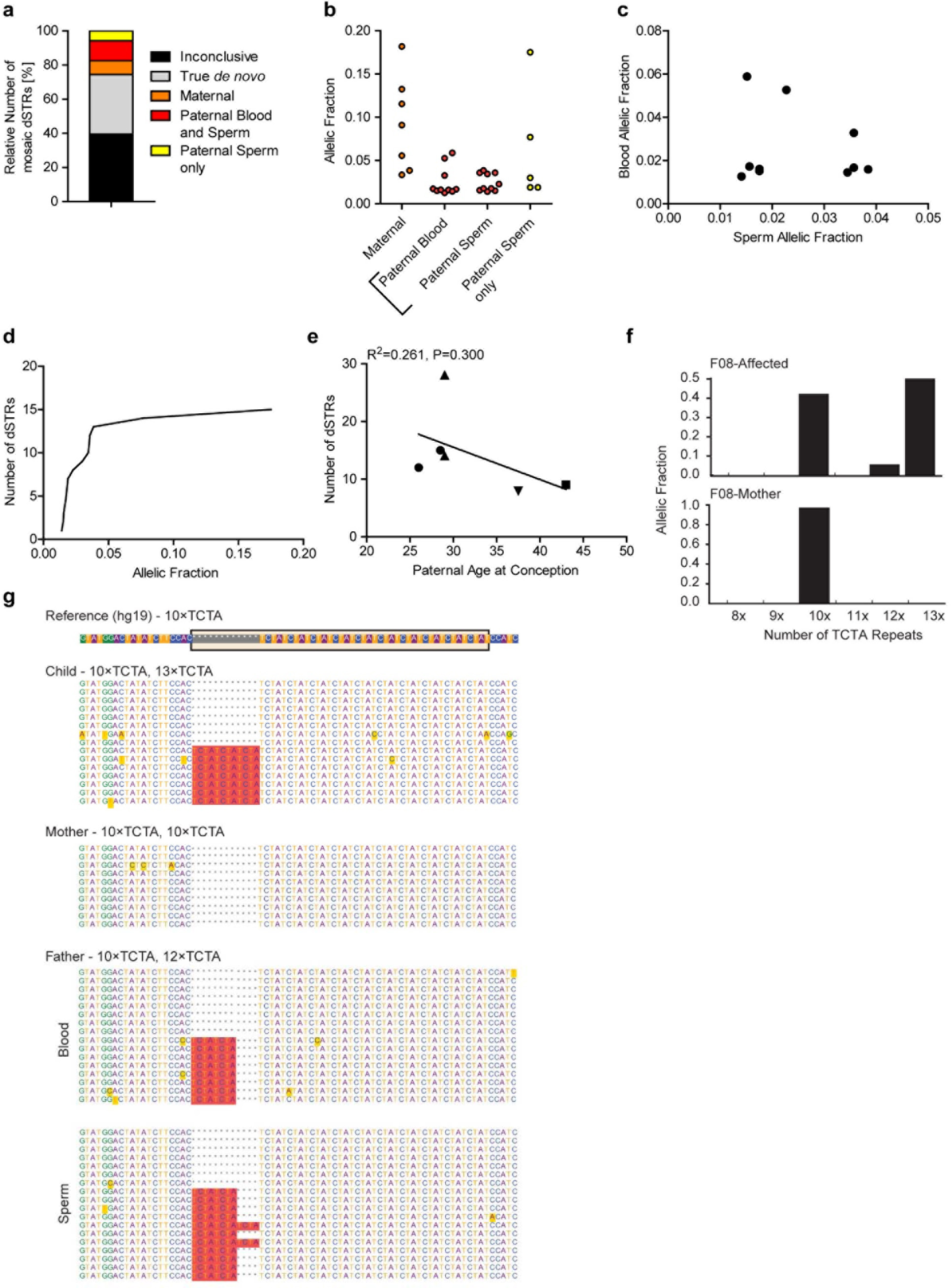
Supplementary graphs for mosaicism detection of dSTRΔs. **a**, Plot showing the relative number of *de novo* short tandem repeat variants (dSTRΔs), whose mosaic status was inconclusive, that were presumed true *de novo* (no evidence outside the child), or mosaic in the mother, the father’s blood and sperm, or the fathers’s sperm only. **b**, Plot of the AF of dSTRΔs that were also found in the mother, the father’s blood and sperm, or the father’s sperm only. **c**, Plot showing the blood and sperm allelic frequencies for those dSTRΔs that were detected in the father. **d**, Plot showing the cumulative relative fraction of mosaic dSTRΔs. **e**, Plot showing the number of mosaic dSTRΔs relative to the paternal age at conception. Line shows a regression curve without positive correlation (R^2^=0.261, P=0.300). Note that this is in disagreement with previous results that showed positive correlation of dSTRΔs. It is most likely a result of the small sample size and stringent requirements for a dSTRΔ to be considered a *de novo* mutation in our data set. **f**, Detailed analysis of the TCTA repeat numbers in the affected’s and maternal blood. **g**, Sample reads showing the presence of a 10x and 13x allele in the child, a homozygous 10x allele in the mother, a 10x and a 12x allele in the father, and the presence of a mosaic 13x allele exclusively in paternal sperm.

